## Supplementary material for "Practical and effective diagnosis of animal anthrax in endemic low-resource settings": S2 File

### S 2 File. Inter-observer agreement

Kappa statistics was implemented using the irr package [1] for R version 3.6.0, and is based on an equation (1) used for two observers or two observations, where $P(a)$ is the proportion of observed agreement by two observers, and $P(e)$ is the probability of agreement due to chance:

$$\mathcal{K} = \frac{P(a) - P(e)}{1 - P(e)} \quad (1)$$

As Kappa values may be affected by the prevalence of anthrax in the samples tested, which might in turn influence the degree of agreement between observers expressed in the proportion of samples determined as positive and negative (i.e. very high or low prevalence would be expected to result in higher bias) [2,3], prevalence- and bias-adjusted Kappa (PABAK) values were computed using formula 2:

$$\mathcal{K}_{PABAK} = 2P(a) - 1 \quad (2)$$

The results for the assessment of agreement in test outcomes of smears stained by one observer and microscopically examined by two observers showed a nearly perfect inter-observer agreement for azure B and PMB, with Kappa scores of 0.94 and 0.95, respectively. However, agreement for Giemsa or Rapi-Diff II stain was moderate with scores of 0.51 and 0.41, respectively. However, when adjusted for prevalence and observer bias, they had substantial agreement, with PABAK values of 0.80 and 0.86.

**Inter-observer agreement for the interpretation of smears stained with different techniques**

| Technique | Number of observations | Cohen's Kappa | Prevalence-bias-adjusted Kappa | z statistic | *p*-value |
|---|---:|---:|---:|---:|---:|
| Azure B | 144 | 0.94 | 0.94 | 11.3 | 0.00 |
| PMB | 84 | 0.95 | 0.95 | 8.71 | 0.00 |
| Giemsa | 140 | 0.51 | 0.80 | 6.01 | 1.91e-09 |
| Rapi-Diff II | 143 | 0.41 | 0.86 | 5.03 | 4.79e-07 |

Inter-observer agreement between two observers on smear samples from the same carcass that were stained separately by the observers using the azure B technique also yielded high agreement, with a Kappa score of 0.94 (z= 7.93, p= 2.22e-15, PABAK = 0.94).

The kappa scores for the agreement between results obtained with the TVLA testing and azure B stain testing was 0.79 ($z = 6.54$, $P > 0.05$). The agreement with PMB was 0.73, ($z = 6.03$, $P > 0.05$) and PABAK scores for both tests yielded the same value as their Kappa scores. Both scores indicate substantial agreement.
