## Supplementary material for "Practical and effective diagnosis of animal anthrax in endemic low-resource settings": S3 Table

**S 3 Table. Primer and probe sequences used in the qPCR reactions, targeting two plasmids and one chromosomal sequence in the *B. anthracis* genome**

| Name of primer/probe | Sequence | Target | Reaction conditions |
|---|---|---|---|
| BA cap fwd[*] TM | GAA GCA GTA GCA CCA GTA AAA CAT C | pXO2 plasmid | 300 nM (0.6 µl of 10µM) |
| BA cap rev[δ] TM | CTT TTA CGT GAC GTC CCA TCA | pXO2 plasmid | 900 nM (1.8 µl of 10µM) |
| BA cap prb[γ] TM | FAM TTG ACG ATG ACG ATG GTT GGT GAC A BHQ1 | pXO2 plasmid | 250 nM (0.5 µl of 10µM) |
| BA lef fwd[*] TM | GGA ACA AAA TAG CAA TGA GGT ACA AGA | pXO1 plasmid | 900 nM (1.8 µl of 10µM) |
| BA lef rev[δ] TM | TTC CGG TGC ATA AAG CTG TAA AAC | pXO1 plasmid | 600 nM (1.2 µl of 10µM) |
| BA lef prb[γ] TM | FAM TTG CAT ATT ATA TCG AGC CAC AGC ATC GTG BHQ1 | pXO1 plasmid | 250 nM (0.5 µl of 10µM) |
| PLF3_f[*] | AAAGCTACAAACTCT GAAATTTGTAAATTG | Chromosomal sequence | 250 nM (0.5 µl of 10µM) |
| PLF3_r[δ] | CAACGATGATTGGA GATAGAGTATTCTTT | Chromosomal sequence | 250 nM (0.5 µl of 10µM) |
| Tqpro_PL3[γ] | FAMAACAGTACGTTT CACTGGAGCAAAAT CAABHQ1 | Chromosomal sequence | 150 nM (0.3 µl of 10µM) |

* Forward primer
δ Reverse primer
γ Probe
