## Supplementary material for "Practical and effective diagnosis of animal anthrax in endemic low-resource settings": S2 Table

**S 2 Table. Comparison of capsule scores obtained with polychrome methylene blue and azure B**

| Capsule strength | Number of samples using azure B stain (n= 102) | Number of samples using polychrome methylene blue stain (n= 102) |
|---|---|---|
| 0 | 40 | 40 |
| +/- | 10 | 9 |
| 1+ | 14 | 23 |
| 2+ | 26 | 20 |
| 3+ | 12 | 10 |
