## Supplementary material for "Practical and effective diagnosis of animal anthrax in endemic low-resource settings": S1 Table

**S 1 Table: Characteristics of the animal carcasses suspected to have died from anthrax, from which samples were collected**

| Variable | Number of samples with data (%) | Number of samples with missing values (%) |
|---|---|---|
| **Species** | **345 (94.0)** | **22 (6.0)** |
| Cattle | 34 (9.3) | |
| Goat | 37 (10.1) | |
| Sheep | 247 (67.3) | |
| Donkey | 16 (4.4) | |
| Giraffe | 3 (0.8) | |
| Antelope | 1 (0.3) | |
| Wildebeest | 2 (0.5) | |
| Zebra | 3 (0.8) | |
| Elephant | 1 (0.3) | |
| **Age** | **330 (96.1)** | **37 (10.1)** |
| Juvenile | 38 (10.4) | |
| Sub-adult | 75 (20.4) | |
| Adult | 217 (59.1) | |
| **Sex** | **172 (46.9)** | **195 (53.1)** |
| Female | 116 (31.6) | |
| Male | 56 (15.3) | |
| **Body condition prior to death (livestock species only, n=335)** | **315 (92.1)** | **11(3.3)** |
| Fat | 162 (48.4) | |
| Normal | 136 (40.6) | |
| Thin | 3 (0.9) | |
| **Intactness of carcasses prior to sampling** | **315 (92.1)** | **52 (14.2)** |
| Intact carcass | 20 (5.4) | |
| Open carcass | 295 (80.4) | |
