## Supplementary material for "Practical and effective diagnosis of animal anthrax in endemic low-resource settings": S2 Fg

**Data informing the latent class model**

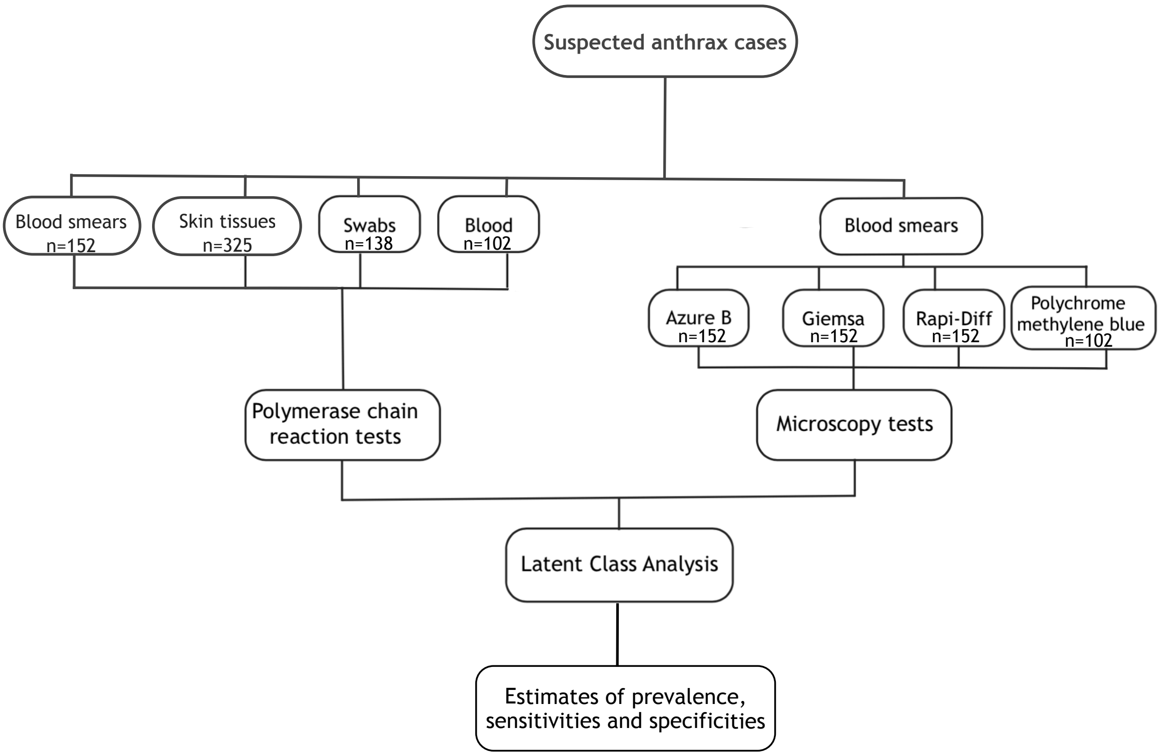


**S 2 Fig. Workflow for samples and data informing the latent class model used to estimate the sensitivities and specificities of different tests used to diagnose suspected anthrax carcasses**
