## Supplementary material for "Practical and effective diagnosis of animal anthrax in endemic low-resource settings": S1 Fig

**S 1 Fig. Chart used to establish presence and strength of *Bacillus anthracis* capsule material**

+/-

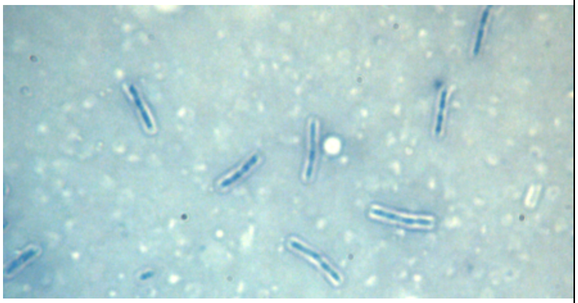

Presence of capsule well demarcated but not metachromatic.

+/-

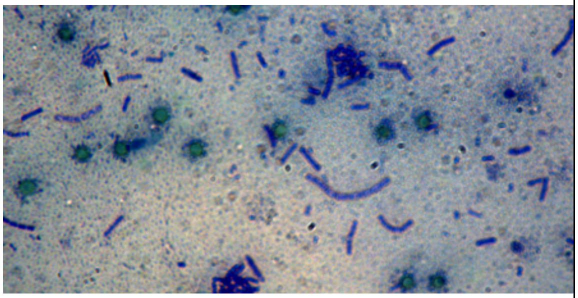

Pink outline faintly visible around the bacilli, but not demarcated.

1+

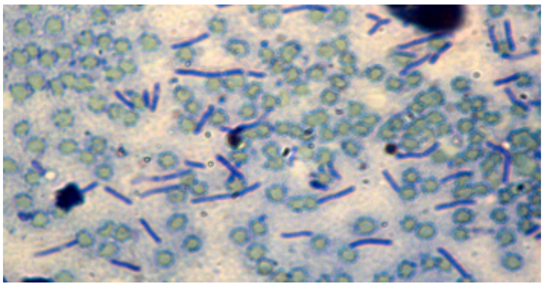

Capsule is visible as a faint pink weakly demarcated around the bacilli.

2+

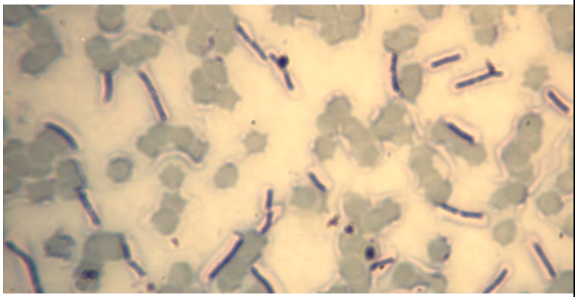

Capsule is moderately stained and well demarcated.

3+

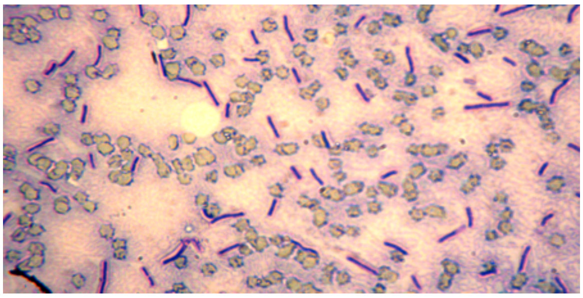

Capsule is strongly stained and well demarcated.

**The strength of the capsule is based on the presence of a demarcated capsule or a 'shadon' surrounding the cell and the metachromatic property of the capsule (Owen *et al.*, 2013). The chart is interpreted subjectively, and all categories (including +/-) are interpreted as *B. anthracis* positive as they show evidence of the presence of a capsule. Samples with a +/- score are so indicated due to the absence of either a demarcated or metachromatic capsule. Images were obtained from pictures of slides examined in the study.**
