## Supplementary material for "Practical and effective diagnosis of animal anthrax in endemic low-resource settings": S1 File

### S 1 File. Stain preparation and staining procedures

#### Azure B staining

Azure B stain was prepared according to the method of Owen *et al.* [1]. Briefly, 0.23% azure B (VWR, United Kingdom) was obtained by dissolving 0.03g of azure B powder in 3ml of 95% ethanol. After gentle swirling for about 30 seconds, 10ml of 0.01% potassium hydroxide solution was added. Air-dried smears were fixed in 99% ethanol for 1 minute and allowed to air-dry. The fixed smears were stained by spreading a drop of azure B solution using a Pasteur pipette and leaving it in contact for approximately 5 minutes. The stain was washed off with water into chlorine solution greater than 10,000 ppm, as part of biosafety measures and to ensure inactivation of any viable *B. anthracis*.

#### Polychrome methylene blue staining

Polychrome methylene blue (PMB) stain (BDH Chemicals, United States of America) was obtained from Public Health England, having aged for more than 12 months and up to 10 years. Staining was carried out according to WHO guidelines [2]. Smears were fixed by dipping in 99% ethanol for 1 minute. Fixed smears were allowed to air dry before adding a drop of PMB stain, which was spread to cover the smear completely. The stain was left on the smear for 30 to 60 seconds and then washed off with water from a wash bottle into chlorine solution greater than 10,000 ppm.

#### Giemsa staining

Giemsa stock solution (0.72% (w/v)) was prepared by dissolving 0.5 g Giemsa stain (Sigma-Aldrich, Germany) in 27 ml glycerol (Sigma-Aldrich, Germany). The solution was heated to 60 °C for 2 hours and then cooled to room temperature, after which 42 ml of methanol was added. The stain was kept in a dark area for more than 3 months to mature. Working solution was prepared by diluting the stock solution 1:20 in phosphate-buffered saline (PBS). Air-dried smears were fixed in absolute methanol for five to seven minutes and left to air dry. The smear was stained for 60 minutes. Stain was washed off with deionised water into chlorine solution greater than 10,000 ppm.

#### Rapi-Diff II staining

Rapi-Diff II stain kit (Vetlab Supplies, United Kingdom) was used as obtained directly from the manufacturer. Reagents included in the kit were methanol-based fixative solution (solution A), eosin Y dye in phosphate buffer (solution B), and polychrome methylene blue in phosphate buffer (solution C). Aliquots of each solution were dispensed into staining containers. Air-dried smears were fixed by dipping slides into solution A for approximately five seconds. The slides were then removed and transferred immediately into solution B, dipping and withdrawing the slides every two seconds for a total of five times. Excess stain was drained with paper towel and the stain was completely rinsed off using PBS from a wash bottle into chlorine solution. Slides were transferred into solution C, dipping and withdrawing the slides five times with two second-intervals in between each dip. Solution C was washed off with PBS into chlorine solution greater than 10,000ppm and the slide allowed to dry.
